# Unveiling Differential Epigenetic Regulation of ABC Transporter Genes in Wild and Domestic Rice in Response to Heavy Metal Stress

**DOI:** 10.1101/2025.11.20.687438

**Authors:** Dipro Sinha, Sougata Bhattacharjee, Bandana Bose, Sananda Mondal

**Affiliations:** IRRI-South Asia Hub, Hyderabad, ICAR-Indian Agricultural Research Institute, New Delhi; Indian Agricultural Research Institute, Hazaribagh, Jharkhand, India; Institute of Agricultural sciences, Banaras Hindu University, Varanasi-221005, UP, India; Department of Crop Physiology, Institute of Agriculture, Visva-Bharati University, Sriniketan-731236, WB, India

**Keywords:** Heavy Metals, Epigenetic Regulation, DNA Methylation, Promoter Analysis, ABC Transporter

## Abstract

Plants are continually subjected to heavy metal (HM) stress, which impairs critical physiological and metabolic processes. ATP-binding cassette (ABC) transporters play an important role in heavy metal detoxification, their epigenetic regulation under heavy metal stress remains poorly understood. Epigenetic changes, such as DNA methylation (5mC) and N6-methyladenine (6mA), are emerging as important regulators of stress-responsive genes. We studied the epigenetic regulation of ABC transporter genes in three *Oryza* species *Oryza sativa indica*, *Oryza rufipogon*, and *Oryza glaberrima* by analysing promoter methylation patterns. Our data show that 5mC is abundant in CHG motifs, whereas 6mA is mostly found at AHCA motifs, indicating sequence-specific methylation preferences across different rice species. Notably, *indica* rice exhibited higher promoter methylation levels compared to *O. rufipogon* and *O. glaberrima*, suggesting a potential link between increased DNA methylation and reduced ABC transporter expression, which may contribute to heightened HM sensitivity in cultivated rice. Phylogenetic analysis further highlighted species-specific variations in methylation patterns, supporting the role of epigenetics in shaping HM stress adaptation. These results provide new insights into the regulatory mechanisms governing heavy metal tolerance in rice and open avenues for epigenetic-based crop improvement strategies.

## Introduction

Plants are constantly exposed to adverse environmental conditions that negatively impact their growth, development, and productivity. Among these stressors, heavy metal (HM) toxicity is a major concern, as excessive metal accumulation disrupts essential physiological and metabolic processes. It is well established that heavy metals such as copper (Cu), iron (Fe), manganese (Mn), zinc (Zn), nickel (Ni), cobalt (Co), cadmium (Cd), and arsenic (As) can be toxic to plants at high concentrations (DalCorso et al., 2008; Hossain et al., 2009; Rascio & Navari-Izzo, 2011). While some metals serve as essential micronutrients, excessive levels can induce oxidative stress, leading to lipid peroxidation, protein oxidation, DNA damage, and enzyme inactivation. To counteract these effects, plants have evolved multiple detoxification mechanisms, including antioxidant defence systems, glyoxalase pathways, and metal sequestration by glutathione (GSH) and metal-binding ligands (Hossain et al. 2012a; Singhal et al. 2023)

Beyond these biochemical defence strategies, epigenetic modifications have emerged as crucial regulators of plant stress responses. DNA methylation, histone modifications, and RNA interference (RNAi) play key roles in fine-tuning gene expression under different environmental stresses (Saraswat et al. 2017). Among the different epigenetic marks, 5-methylcytosine (5mC) is one of the most extensively studied DNA modifications in plants, found at CG, CHG, and CHH sites (H = A, T, or C). This modification regulates gene activity by reshaping chromatin organization and thereby modulating the accessibility of regulatory regions. In plants, DNA methylation is maintained through the coordinated action of MET1 (responsible for CG methylation, homologous to animal methyltransferases), CMT3 (a plant-specific enzyme that preserves CHG methylation), and the RNA-directed DNA methylation (RdDM) pathway, which establishes de novo CHH methylation (Du et al. 2012; Matzke et al. 2025; Woo et al. 2008). Due to its dynamic and reversible nature, DNA methylation serves as an adaptive mechanism that enables fine-tuned regulation of stress-responsive genes (Campbell & Kleckner 1990; Feng & Jacobsen 2011; Flusberg et al. 2010).

Another key epigenetic modification, N6-methyladenine (6mA), has recently been recognized as an important regulatory mark in plants. Unlike 5-methylcytosine (5mC), which primarily represses gene expression, 6mA has been linked to both gene activation and repression, depending on its genomic context. Studies suggest that 6mA plays a crucial role in transcriptional regulation, chromatin accessibility, and response to environmental stresses (Hossain et al. 2012b; Pukkila et al. 1983; Zhou et al. 2018). However, the functional role of 6mA methylation in stress-responsive genes, particularly in rice under heavy metal stress, remains largely unexplored.

A major gene family involved in heavy metal stress response is the ATP-binding cassette (ABC) transporter family. ABC transporters constitute one of the largest protein families, facilitating the transport of diverse substrates, including inorganic ions, phytohormones, secondary metabolites, and heavy metals (Locher, 2008). These transporters function as ATP-driven pumps, playing a crucial role in metal homeostasis, detoxification, and sequestration (Taiz et al. 2015). For example, specific ABC transporters mediate the vacuolar sequestration of flavonoids and anthocyanins, while others regulate the transmembrane transport of the stress hormone abscisic acid (ABA) (Espinola et al. 2025; Hossain et al. 2010). Given their essential role in cellular transport and stress adaptation, the regulation of ABC transporter genes is critical for plant responses to heavy metal toxicity (Saraswat et al. 2017; Vanyushin & Ashapkin, 2011).

In this study, we analyzed the candidate promoter regions of ABC transporter genes to investigate the role of DNA methylation (5mC) and N6-methyladenine (6mA) in regulating heavy metal tolerance across three *Oryza* species—*Oryza sativa indica*, *Oryza rufipogon*, and *Oryza glaberrima*. Our sequence enrichment analysis revealed that 5mC sites are enriched in CHG motifs, whereas 6mA sites predominantly occur at AHCA motifs, indicating sequence-specific preferences for methylation (Villiers et al. 2011). Notably, we observed higher methylation levels in the promoter regions of ABC transporter genes in *indica* rice compared to *O. rufipogon* and *O. glaberrima*. Since promoter methylation is often linked to transcriptional repression, these findings suggest that differential epigenetic regulation may influence ABC transporter expression and heavy metal tolerance among rice species.

By elucidating the methylation patterns of both 5mC and 6mA in ABC transporter genes, this study provides new insights into the epigenetic mechanisms underlying heavy metal stress responses in rice. Future research integrating transcriptomic and functional analyses will be crucial for understanding how these epigenetic modifications impact gene expression and stress adaptation. Ultimately, modulating epigenetic marks could serve as a potential strategy for enhancing heavy metal tolerance in rice.

## 2. Materials and Methods

### 2.1 Data Collection

The amino acid sequence of ATP-binding cassette (ABC) transporter (Stroud et al. 2013; Zemach et al. 2013) proteins was retrieved from *Arabidopsis thaliana* (NP_181228.1 ABCB1; Gene ID=818265) database, and it was used as a query sequence. Homology was established using HMM profiles (PF00005, threshold E-value 1.0), validated by Pfam and CDS searches, and cross-checked with BLASTp at E-value ≤ 10[[. Three rice species were considered for analysis, and searched for the homologs of the above mentioned gene *viz. O. sativa indica* (https://plants.ensembl.org/Oryza_indica/Info/Index)*, O. rufipogon* (https://plants.ensembl.org/Oryza_rufipogon/Info/Index) and *O. glaberrima* (https://plants.ensembl.org/Oryza_glaberrima/Info/Index).

### 2.2 Genome-Wide Identification

The HMM (Hidden Markov Model) profiles of the ABC transporter domain were obtained from the Pfam database (http://www.pfam.xfam.org/). Initial identification of the ABC transporter domain-containing genes in the genome and protein sequence databases was performed by scanning the entire HMM profiles with the conserved sequences of the ABC transporter domain (PF00005) as a query at a threshold E-value of 1.0. After establishing an initial shortlist of candidates based on HMM profiles, the putative ABC transporter gene sequences were subjected to Pfam and CDS searches to validate the ABC transporter domains. Because the HMM search could fail in identifying truncated ABC transporter domains, an additional BLASTp search was also performed to reveal the signatures of partial domains among the putative ABC transporter-encoding genes. For the BLASTp search, the confirmed ABC transporter-encoding genes identified by HMM profiles were used as queries against the entire protein sequence database of the three at a threshold e-value of 10^−5^.

### 2.3 Species selection

Upstream candidate promoter regions (2 kb) of ABC transporter genes in three *Oryza* species *O. sativa indica*, *O. glaberrima*, and *O. rufipogon* were selected for this study. The selection of these species was based on their phylogenetic relationships, with *O. sativa indica* representing a highly cultivated variety, *O. glaberrima* serving as an African domesticated rice species, and *O. rufipogon* acting as the wild progenitor. This selection allows for a comparative analysis of epigenetic modifications in domesticated vs. wild species and their potential impact on gene regulation.

### 2.4 Extraction of Promoter Region

Promoter regions of ABC transporter genes were extracted from the Ensembl Plants database using the Biomart tool (http://plants.ensembl.org/info/data/biomart/index.html). The sequences were obtained from their respective genome assemblies: *O. sativa indica* (ASM465v1), *O. glaberrima* (Oryza_glaberrima_v1), and *O. rufipogon* (OR_W1943). A 2-kb upstream region from the transcription start site (TSS) was extracted for each gene to ensure the inclusion of potential regulatory elements, such as transcription factor binding sites and DNA methylation sites. The extracted promoter sequences were curated to remove ambiguous bases, ensuring high-quality input data for downstream methylation analysis and comparative promoter studies across the three *Oryza* species.

The 2 kb upstream regions of identified ABC transporter genes were extracted from annotated genome sequences of the three species using publicly available databases and in-house bioinformatics pipelines. These sequences were carefully selected to ensure high sequencing coverage and accurate promoter annotation. The region was chosen to capture potential regulatory elements, such as transcription factor binding sites and DNA methylation marks, which may influence the expression of ABC transporter genes.

A comparative promoter analysis was performed to investigate the distribution and conservation of 5mC and 6mA methylation sites across these species. This approach helps in understanding how methylation patterns have evolved during rice domestication and how they contribute to differential gene regulation in various environmental conditions. The findings from this analysis will provide key insights into epigenetic modifications governing ABC transporter activity, which is crucial for plant responses to nutrient transport, stress resistance, and metabolic functions in rice.

### 2.5 Pre-processing of Sequences

The extracted 2-kb upstream sequences of ABC transporter genes from all three species were pre-processed to ensure high-quality input for methylation site prediction. Any ambiguous bases (containing “N”) were removed using the SeqKit tool (https://bioinf.shenwei.me/seqkit/). After filtering, the promoter sequences were fragmented into 41-bp overlapping segments using the "split fasta" function of the Sequence Manipulation Suite (https://www.bioinformatics.org/sms2/split_fasta.html). The processed FASTA file was then used as input for the GB5mCPred and MethSemble-6mA servers to predict 5mC and 6mA methylation sites, respectively. These predicted methylation sites were further analyzed to identify conserved and species-specific epigenetic modifications within the promoter regions of ABC transporter genes.

### 2.6 Prediction of 6mA Sites in the Promoter of ABC Transporter Genes

For the prediction of 6mA sites in the upstream regions of ABC transporter genes, a single-species model approach was applied using *O. sativa* as the reference (Sinhaspecies (Sinha et al., 2023). The MethSemble-6mA server was employed to predict 6mA methylation sites across the 2-kb promoter regions of *Oryza sativa indica*, *O. glaberrima*, and *O. rufipogon*. This approach ensures that the model, trained specifically on Poaceae species, provides high-accuracy predictions tailored to rice and its wild relatives. By focusing on a single reference species, this methodology reduces species bias while maintaining high sensitivity and specificity in detecting methylation patterns that may regulate ABC transporter gene expression. The predicted 6mA sites were further analyzed to identify conserved and species-specific methylation patterns, providing insights into the epigenetic regulation of transporter genes in different *Oryza* species.

### 2.7 Identification of 5mC Sites in the Promoter Region of the ABC Transporter Genes

To predict 5-methylcytosine (5mC) sites in the promoter region of the ABC transporter domain in *O. sativa indica*, *O. glaberrima*, and *O. rufipogon*, the GB5mCPred tool has been used. The promoter regions (2 kb upstream of the transcription start site) will be extracted from genome annotation files using ENSEMBL Plants Biomart or in-house scripts. These sequences will then be converted into numerical feature vectors using Di-nucleotide Frequency (DNF), Mono-nucleotide Binary Encoding (MBE), and Chemical Properties of Nucleotides (NCP). The prediction will be performed using Random Forest, Gradient Boosting, and Support Vector Machine (SVM) models, as these have been found to be the most accurate for 5mC site identification (Sinha et al., 2024). The predictions can be executed via the GB5mCPred web server (http://cabgrid.res.in:5474/). The identified 5mC sites in the promoter region will be analyzed for conservation across the three *Oryza* species, and their potential regulatory impact on ABC transporter gene expression will be assessed.

### 2.8 Mapping of 6mA Sites and Phylogenetic Analysis

The predicted 6mA sites in the 2-kb promoter regions of ABC transporter genes were analyzed to identify regions with the highest methylation density. These sites were mapped using the MapChart tool to visualize their distribution across the promoter sequences. To assess evolutionary conservation, a standalone BLASTn search was performed using the NCBI BLAST+ tool, with *O. sativa indica* ABC transporter gene promoters as queries and *O. glaberrima* and *O. rufipogon* promoter sequences as the database. Hits with 100% sequence identity were retained for multiple sequence alignment (MSA), which was performed using ClustalW.

To understand the evolutionary relationships of methylated promoter sequences, phylogenetic analysis was conducted using the Maximum Likelihood Method (MLM) in MEGA XI (Tamura et al., 2021). A Newick tree file was generated and visualized using the iTOL tool (https://itol.embl.de/). This analysis helped in identifying conserved and species-specific 6mA methylation patterns, providing insights into the evolutionary role of DNA methylation in regulating ABC transporter genes across different *Oryza* species.

### 2.9 Motif Enrichment Analysis of Methylated Promoter Regions

To investigate potential regulatory elements associated with DNA methylation, motif enrichment analysis was performed using TBtools (Chen et al. 2020). The 5mC-enriched sequences and their non-methylated counterparts, as well as the 6mA-enriched sequences and non-methylated sequences, were analysed separately. This approach allowed the identification of overrepresented motifs that may play a role in transcriptional regulation of ABC transporter genes.

The enriched motifs were compared across *O. sativa indica*, *O. glaberrima*, and *O. rufipogon* to determine conserved regulatory elements associated with 5mC and 6mA modifications. Differences in motif occurrence between methylated and non-methylated sequences provided insights into the potential influence of epigenetic modifications on transcription factor binding. Additionally, species-specific motif variations were examined to understand how methylation-associated regulatory elements have evolved across domesticated and wild *Oryza* species. This analysis contributes to a deeper understanding of methylation-mediated gene regulation in ABC transporter genes, highlighting key sequences that may influence gene expression, stress response, and metabolic functions in rice.

## 3. Results

### 3.1. Putative ABC Transporter Domain Identification and Promoter Extraction

HMM search was performed to identify ABC transporter genes across three *Oryza* species—*O. sativa indica*, *O. glaberrima*, and *O. rufipogon*. Initially, a larger set of genes was identified; however, after applying stringent filtering criteria, only 31 high-confidence genes containing the ABC transporter domain were retained. These included 18 genes from O. sativa indica, 5 from *O. glaberrima*, and 8 from *O. rufipogon*.

The filtering process involved removing redundant sequences and low-confidence hits (as described in section 2.1), ensuring that only genes with significant HMM scores and complete domain structures were selected. A comparative analysis of these genes across the three species revealed both conserved and species-specific ABC transporter genes, providing insights into the evolutionary adaptation and functional diversification of this gene family in domesticated and wild rice species. This refined dataset serves as the basis for further analysis of regulatory elements, methylation patterns, and transcriptional control mechanisms in *Oryza* species.

To explore the regulatory mechanisms controlling the expression of these 31 selected ABC transporter genes, their 2-kb upstream promoter regions were retrieved from the Ensembl Plants Biomart database. After filtering and ensuring sequence quality and completeness, a total of 23 promoter sequences were successfully extracted. Some genes did not yield retrievable promoters, likely due to incomplete genome annotations or missing upstream sequences in the database. These promoter sequences provide a comprehensive dataset for regulatory analysis, allowing for the identification of potential transcription factor binding sites, DNA methylation patterns, and conserved cis-regulatory elements. This dataset forms the foundation for further 5mC and 6mA methylation site prediction, motif enrichment analysis, and comparative regulatory studies, helping to uncover epigenetic modifications that influence ABC transporter gene expression in *Oryza* species.

### 3.2. Study on Methylation Pattern

The identification of N6-methyladenine (6mA) (Suppl. Material 1) and 5-methylcytosine (5mC) (Suppl. Material 2) methylation sites across 23 promoter regions revealed distinct methylation patterns, with varying levels of modification observed across different genes and species. Some genes exhibited extensive methylation across multiple positions (e.g., BGIOSGA014966, with 11 methylation sites for 6mA, and BGIOSGA005326, with 8 methylation sites for 5mC), while others had only one or two detected modifications. The presence of NA values indicates the absence of methylation at certain loci.

#### 3.2.1 Species-Specific Methylation Variations

Species-specific differences were evident for both methylation types. The average 6mA methylation rates were 5.5 in *O. indica*, 5.2 in *O. glaberrima*, and 5.1 in *O. rufipogon*, while 5mC methylation rates were 3.6 in *O. indica*, 3.0 in *O. glaberrima*, and 3.16 in *O. rufipogon*. These differences suggest distinct epigenetic regulatory mechanisms that may influence gene expression and species adaptation.

#### 3.2.2 Functional Implications and Gene Regulation

The distribution of 6mA and 5mC sites across promoter regions suggests potential roles in gene regulation, chromatin accessibility, and transcriptional activity. Genes such as BGIOSGA006985 and ORUFI02G39650 exhibited eight 6mA methylation sites, indicating potential regulatory significance, while genes like BGIOSGA023241 and ORUFI08G26390 (annotations provided in suppl. file 3 from suppl. figure 1- suppl. figure 4) showed minimal modifications, with only two or three methylation sites. From a regulatory perspective, 6mA methylation has been linked to both transcriptional activation and repression, depending on the genomic context, whereas 5mC methylation is often associated with chromatin remodelling, where hypermethylation in promoters contributes to transcriptional repression and hypomethylated promoters are linked to active gene expression (**Fig. 1A-1F**).

**Figure 1.**
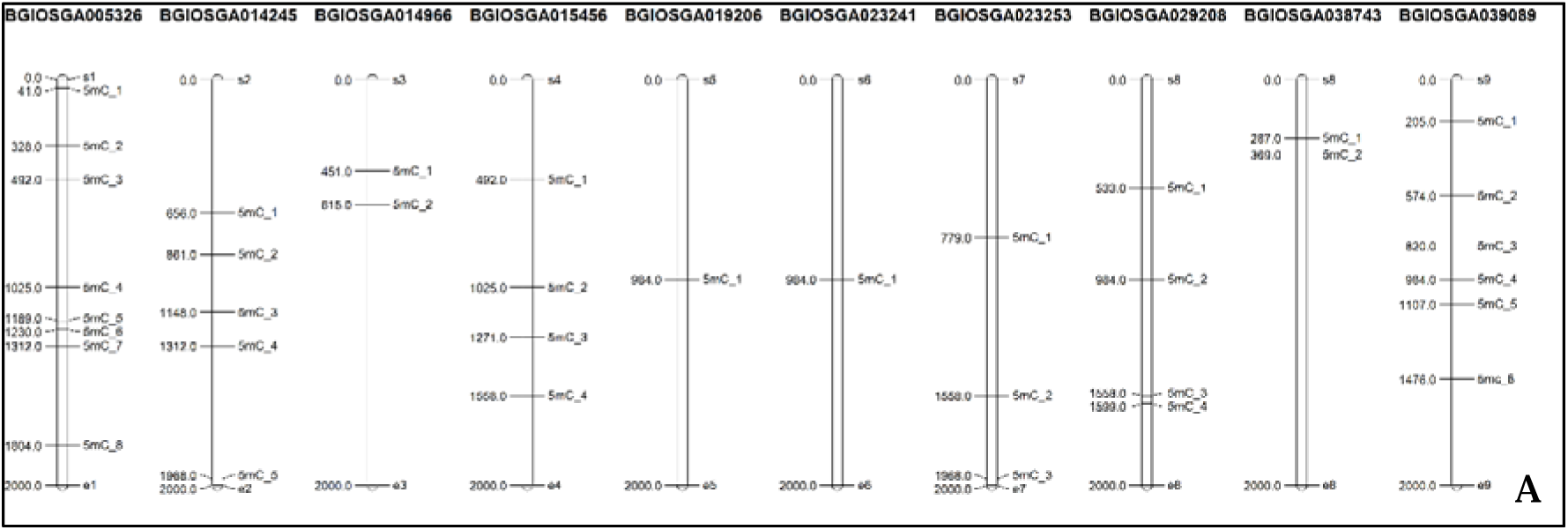
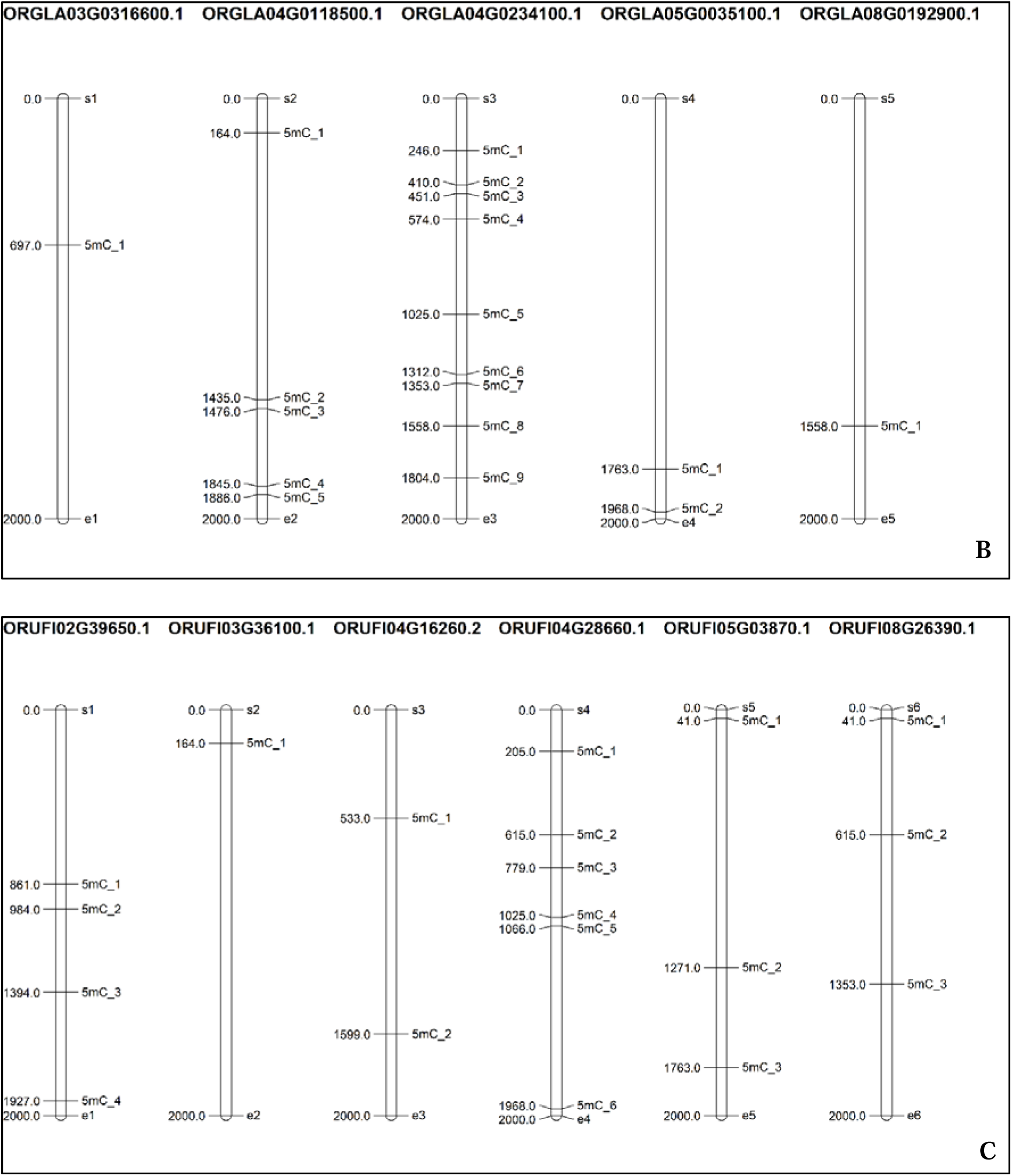
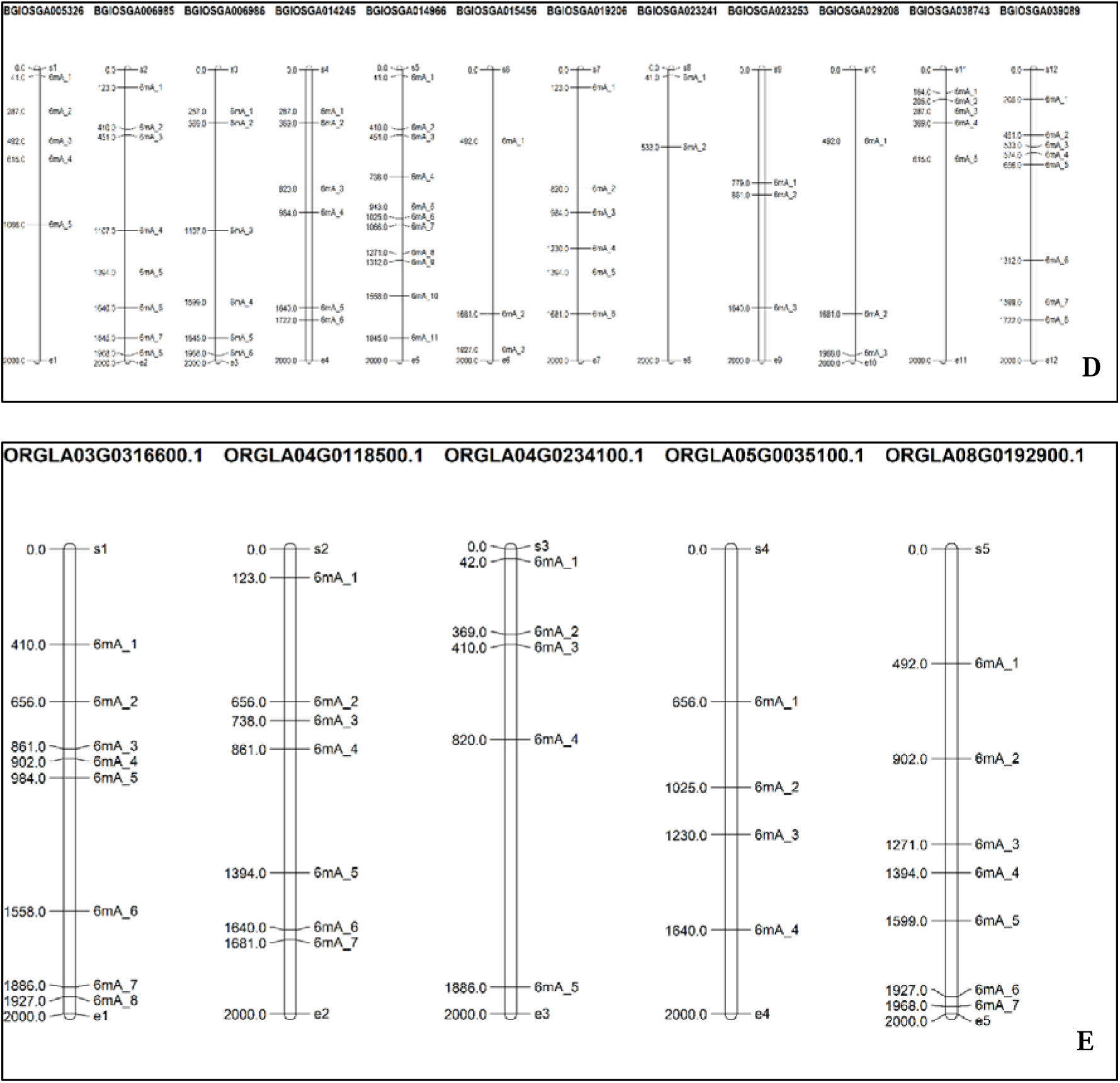
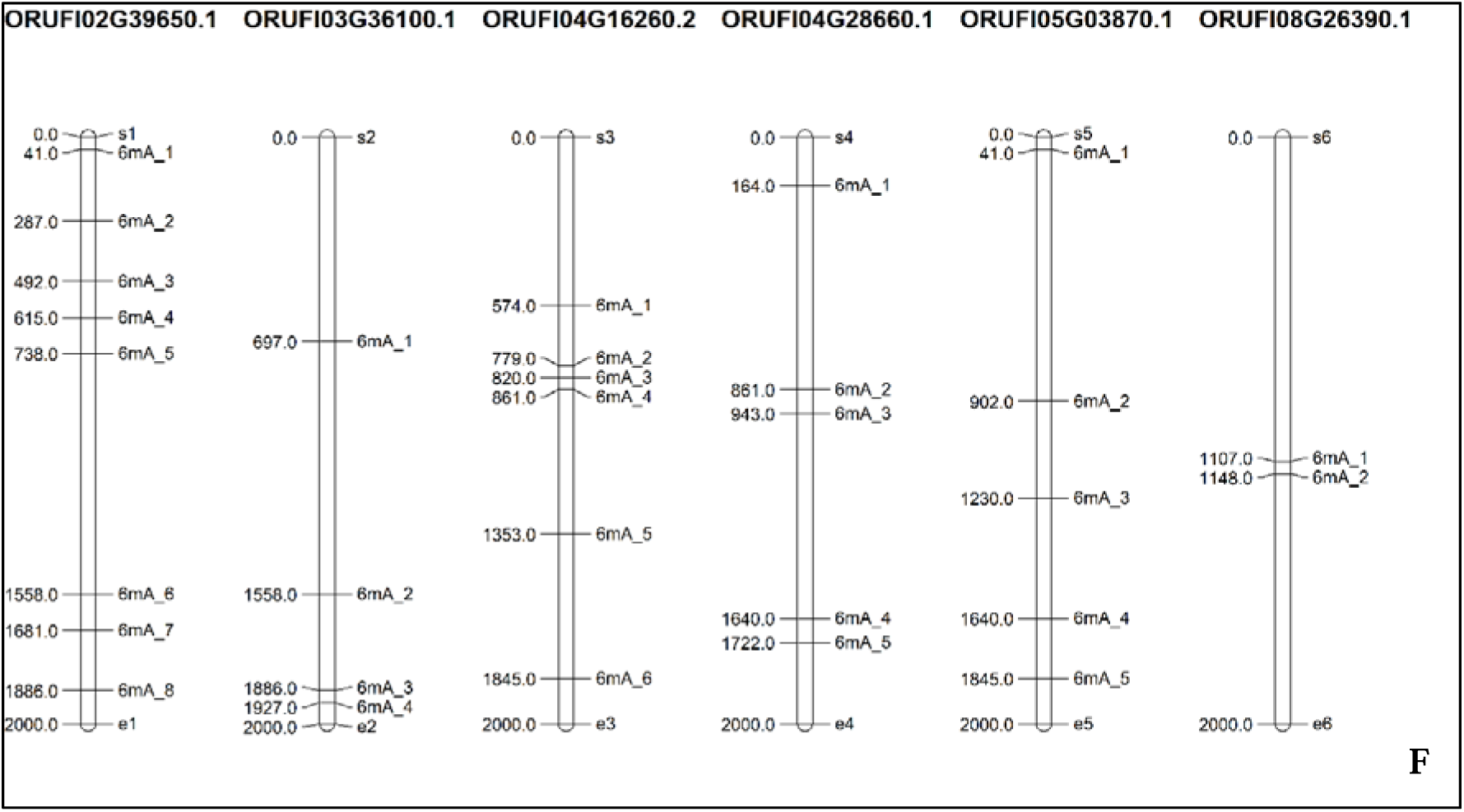
These figures illustrate the methylation pattern (both 5mC and 6mA) in three *Oryza* Species – **A**. 5mC distribution in *O. sativa indica,* **B.** 5mC distribution in *O. glaberrima,* **C.** 5mC distribution in *O. rufipogon,* **D**. 6mA distribution in *O. sativa indica,* **E**. 6mA Distribution in *O. glaberrima,* **F.** 6mA Distribution in *O. rufipogon*

### 3.3. Phylogenetic Analysis

The shortlisted genes with identical sequences containing an ABC transporter domain, identified using BLASTn search, along with their corresponding promoter regions. To understand the evolutionary conservation of 6mA and 5mC methylation patterns across identified candidate ABC transporter genes, we generated two phylogenetic trees—one for the genes and another for the promoter regions (**Fig. 2A and 2B**).

**Figure 2.**
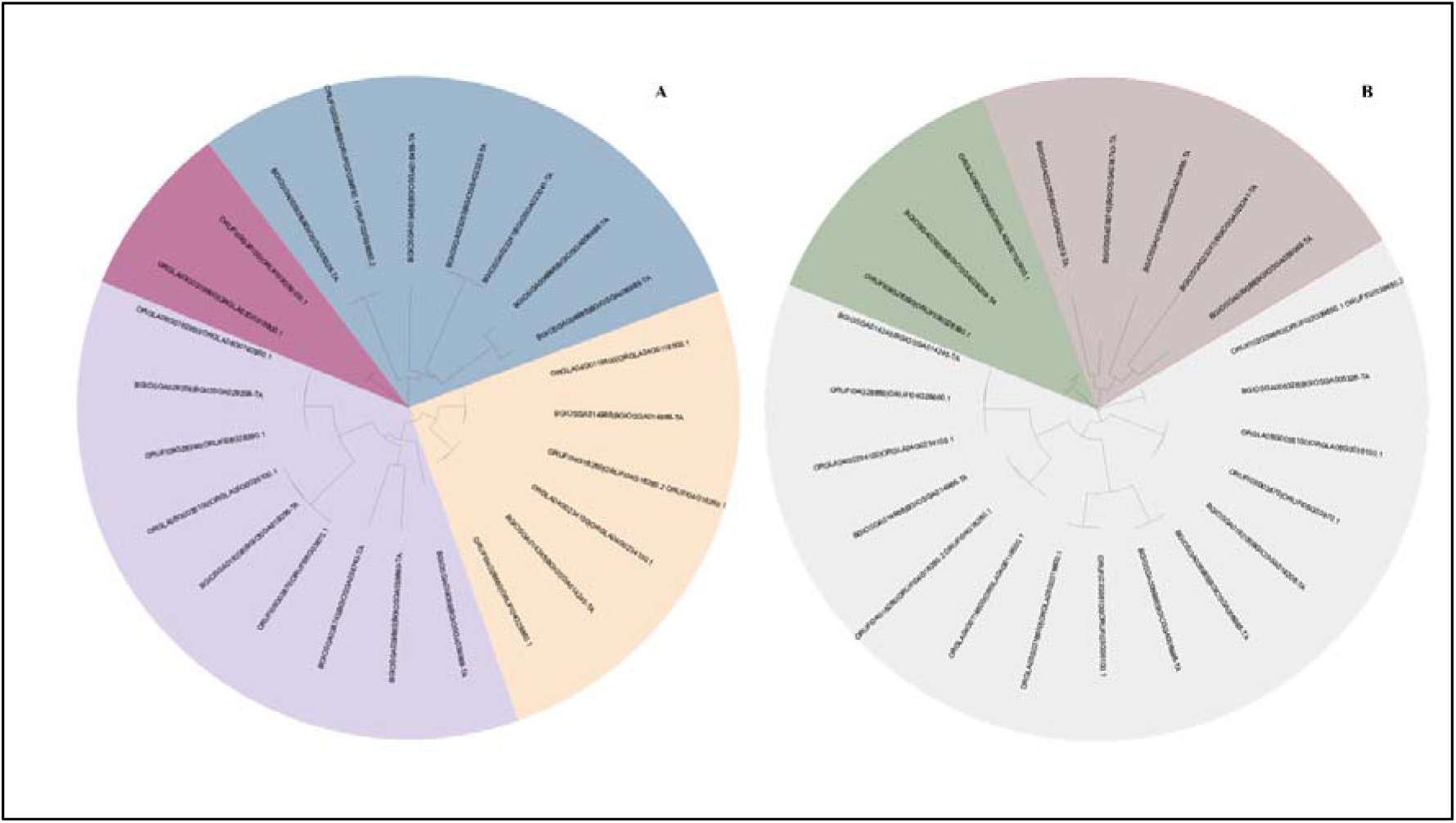
Phylogenetic analysis of three *Oryza* species – **A**. Phylogenetic analysis of genes containing the ABC transporter domain **B.** Phylogenetic analysis of promoter region genes containing the ABC transporter domain

The gene phylogenetic tree was classified into four clades, while the promoter tree was divided into five clades. We analyzed the conservation of 6mA and 5mC methylation sites across these clades. Among the studied species, *O. rufipogon* exhibited the least commonality with other rice species.

For further analysis, we selected representative genes from different clades to investigate the 6mA and 5mC methylation positions in greater detail. The findings from this analysis are elaborated in the Discussion section.

### 3.4. Motif Analysis

Sequence enrichment analysis visually represents nucleotide frequency and conservation across aligned sequences, comparing methylated and non-methylated sites for 5-methylcytosine (5mC) and 6-methyladenine (6mA) (**Fig. 3A-D**). In our sequence enrichment analysis, we examined the sequence composition surrounding putative methylation sites to identify conserved motifs associated with DNA methylation. Our findings reveal that sequences containing putative 5mC sites predominantly feature the CHG motif, where H represents A, T, or C, suggesting a preference for methylation at CHG sites. This observation aligns with known plant and fungal methylation patterns, where cytosine methylation frequently occurs in the CHG context, playing a critical role in gene regulation and epigenetic silencing. Additionally, sequences associated with putative 6mA sites were found to be enriched with the AHCA motif, where H can be A, T, or C, indicating a potential sequence specificity for adenine methylation. The presence of this conserved motif suggests that 6mA methylation may preferentially occur in distinct sequence contexts, potentially influencing transcriptional regulation. These findings provide valuable insights into the sequence determinants of DNA methylation and their potential functional roles in genome regulation. This analysis was conducted on three *Oryza* species: *O. sativa indica*, *O. glaberrima*, and *O. rufipogon*, allowing us to explore species-specific variations in methylation patterns and sequence motif conservation.

**Figure 3.**
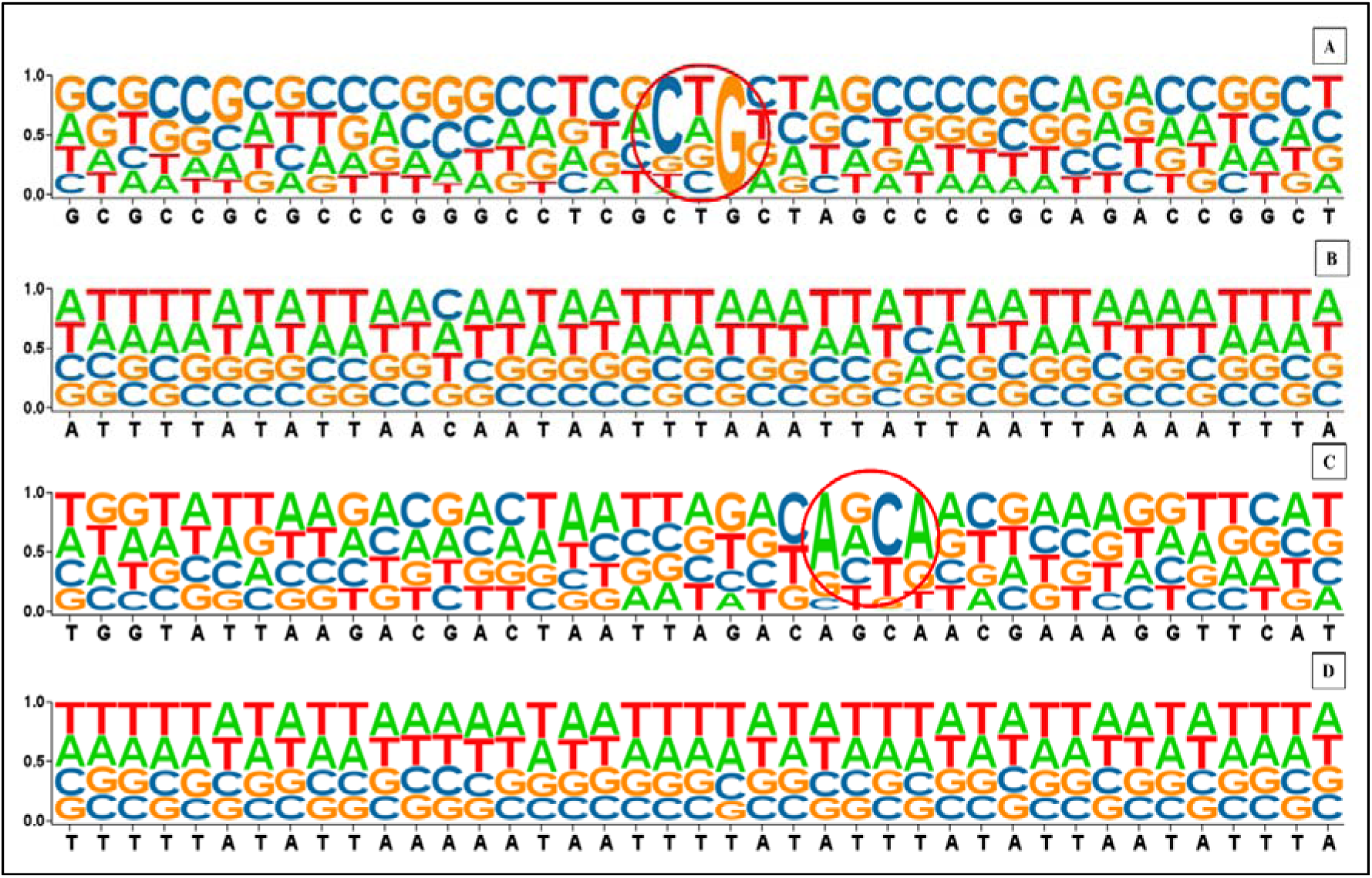
Motif enrichment analysis of sequences associated with DNA methylation. **A**. Sequence containing 5mC sites, **B**. Sequences without 5mC sites C. Sequence containing 6mA sites D. Sequences without 6mA sites

## 4. Discussion

Heavy metal (HM) stress is a significant environmental factor that affects plant growth and development by disrupting essential physiological and metabolic processes (Singhal et al., 2023). The role of 6mA methylation in stress tolerance was earlier reported comprehensively (Zhang et al. 2018). In this study, we investigated the epigenetic regulation of heavy metal tolerance in three *Oryza* species *O. sativa indica*, *O. rufipogon*, and *O. glaberrima*. Our findings indicate that indica rice is more sensitive to heavy metal stress compared to *O. rufipogon* and *O. glaberrima*, suggesting that differences in genetic and epigenetic regulation contribute to species-specific variations in HM stress response mechanisms (Cao & Jacobsen, 2002; Mayaba et al 2020).

One of the key observations from our study is the differential methylation patterns in the promoter regions of the ABC transporter domain, which is reported to play a critical role in metal ion transport and detoxification (Taiz et al. 2015; Espinola et al., 2025)). Our results show that *indica* rice exhibits a higher methylation rate in the promoter regions of ABC transporter genes compared to the other two species. Since promoter methylation is known to regulate gene expression by influencing chromatin accessibility (Zhou et al. 2018), increased methylation in *indica* rice may be associated with transcriptional repression of ABC transporter genes. This could limit the plant’s ability to manage metal toxicity effectively, leading to increased sensitivity to heavy metal stress. In contrast, *O. rufipogon* and *O. glaberrima*, which have lower promoter methylation in these genes, may maintain higher ABC transporter activity, allowing for better metal detoxification and stress adaptation.

To further understand the sequence-level features influencing methylation, we performed sequence enrichment analysis, which allowed us to identify conserved motifs associated with putative 5-methylcytosine (5mC) and 6-methyladenine (6mA) sites. Our findings reveal that 5mC methylation is predominantly enriched in CHG motifs (where H = A, T, or C) (Sinha et al., 2024), which is consistent with known plant methylation patterns, particularly in gene body regions and transposable elements. CHG methylation is often associated with epigenetic silencing, which may play a role in regulating ABC transporter gene expression under stress conditions. Similarly, we found that 6mA methylation sites are enriched in the AHCA motif (where H = A, T, or C), suggesting a potential sequence preference for adenine methylation. Given that 6mA methylation has been linked to transcriptional regulation (Sinha et al. 2023), its presence in conserved motifs within ABC transporter promoter regions suggests that this modification may influence gene activation or repression in response to environmental stimuli.

The identification of species-specific differences in methylation patterns and motif conservation highlights the role of epigenetic regulation in heavy metal tolerance. Since DNA methylation is a reversible modification, it provides plants with a dynamic mechanism to fine-tune gene expression in response to environmental stressors. The higher promoter methylation in indica rice may contribute to its increased sensitivity to heavy metal stress, whereas the relatively lower methylation levels in *O. rufipogon* and *O. glaberrima* may facilitate greater stress resilience by maintaining active ABC transporter gene expression.

Future studies should focus on integrating transcriptomic and functional validation approaches to establish a direct link between DNA methylation and ABC transporter gene expression under heavy metal stress conditions. Additionally, exploring methylation-associated transcription factor binding could provide further insights into the regulatory networks controlling stress responses in rice. These findings have important implications for rice breeding and genetic engineering, as modulating epigenetic marks could be a potential strategy to enhance heavy metal tolerance in rice varieties.

Overall, our study underscores the significance of DNA methylation in regulating HM stress-responsive genes and provides new insights into the interplay between epigenetics and environmental adaptation in rice. By elucidating the sequence motifs and methylation patterns associated with heavy metal tolerance, our findings contribute to a deeper understanding of how epigenetic regulation can be leveraged for crop improvement.

## 5. Conclusion

This study highlights the role of epigenetic regulation in heavy metal tolerance across three *Oryza* species—*O. sativa* indica, *O. rufipogon*, and *O. glaberrima*. Our findings indicate that indica rice exhibits higher sensitivity to heavy metal stress, which correlates with increased DNA methylation in the promoter regions of ABC transporter genes. Since ABC transporters play a key role in metal ion transport and detoxification, their epigenetic regulation may directly influence the plant’s ability to tolerate heavy metals. Additionally, our sequence enrichment analysis revealed that 5mC sites are enriched in CHG motifs, while 6mA sites predominantly occur in AHCA motifs, suggesting sequence-specific methylation preferences. The species-specific differences in methylation patterns indicate that epigenetic modifications serve as a regulatory mechanism shaping stress adaptation in rice. Future studies should focus on integrating transcriptomic and functional analyses to further elucidate the impact of methylation on gene expression and stress resilience. These insights pave the way for utilizing epigenetic modifications as potential targets for breeding or engineering rice varieties with enhanced heavy metal tolerance.

## Supplementary material

Supplementary material is available on the publisher’s website along with the published article.

## Authors Contribution

DS designed the experiments. DS, and SB performed the experiments and processed raw data. DS and SM designed the figures. DS, SB, SM and BB contributed to the writing of the manuscript. All the authors read and approved the final version of the manuscript.

## Data Availability

All the data and supportive information are available within the article.

## Ethics approval and consent to participate

Not applicable.

## Human and animal rights

No animals/humans were used for studies that are basis of this research

